# Zika virus sequence and secondary structure analysis using VADR

**DOI:** 10.64898/2026.09.21.753218

**Authors:** Emma B. Dickinson, Eric P. Nawrocki

## Abstract

Zika virus is a mosquito-borne flavivirus that caused a major epidemic in the Americas in 2015 and 2016 and is associated with congenital birth defects and neurological disease. Like other flaviviruses, it depends on conserved sequence and RNA structural elements that carry out essential steps in its life cycle and in evading host immune defenses. Public sequence databases and annotation tools give free access to Zika sequence data, but comparable structural information is sparse. We built a computational model of Zika sequence and secondary structure for use with the VADR software package, which validates and annotates Zika sequences for GenBank submission and assigns each sequence to a genetic lineage following a published classification scheme. On a test set of 1254 Zika sequences, 97.2% passed validation, and the model’s annotations agreed with those of an existing annotation tool (VIGOR4) for 99.3% of the sequences compared. We also extended VADR to draw secondary structure diagrams of each sequence’s structural elements in its linear and circular conformations using R2DT, and we submitted a newly characterized Zika pseudoknotted RNA element to the Rfam database. Together these freely available resources provide secondary structure information for every Zika sequence.

## INTRODUCTION

Zika virus is a positive-sense single-stranded RNA virus of the family *Flaviviridae*, genus *Flavivirus* (renamed *Orthoflavivirus* by the International Committee on Taxonomy of Viruses in 2023 (1)). First isolated from a rhesus macaque in the Zika Forest of Uganda in 1947 (2), it caused only sporadic human infections across Africa and Asia for decades before it began to drive large epidemics. An outbreak on Yap Island in 2007 infected an estimated 5000 people (73% of residents) (3), and a larger epidemic followed in French Polynesia in 2013 and 2014 (4). The virus then reached the Americas in 2015 and 2016, where an explosive epidemic centered in Brazil revealed the severity of congenital Zika disease. Infection during pregnancy causes microcephaly and other congenital brain defects (5), and Brazil alone recorded 1950 confirmed cases of infection-related microcephaly over these two years (6). Infection in adults has also been linked to Guillain-Barré syndrome (7), and in February 2016 the World Health Organization declared the epidemic a Public Health Emergency of International Concern (8). Zika virus is transmitted mainly by *Aedes aegypti* mosquitoes, but also through sexual contact and from mother to fetus, and it has now been reported in more than 90 countries and territories (8). The virus comprises two lineages, African and Asian, with the Asian lineage responsible for the Pacific and American outbreaks (9, 10). Seabra et al. divide the Asian lineage further, into an Asiatic and an American and Oceania sublineage, giving the three clades ZA, ZB.1 and ZB.2 that we use throughout.

The Zika virus genome is approximately 10.8 kb (10,808 nucleotides in the reference strain NC 035889). It contains a single open reading frame that encodes one polyprotein of roughly 3400 amino acids. Host and viral proteases cleave this polyprotein co- and post-translationally into three structural proteins, the capsid (C), precursor membrane (prM), and envelope (E), and seven nonstructural proteins, NS1, NS2A, NS2B, NS3, NS4A, NS4B, and NS5 (11). The host signal peptidase acts on the lumenal side of the endoplasmic reticulum membrane and the viral NS2B-NS3 protease on the cytoplasmic side, while host furin later cleaves prM into a pr peptide and mature M peptide. Counting these precursor and intermediate products, including the anchored and mature forms of C and the small 2K peptide, the polyprotein is the source of 14 mature peptides in total.

Beyond encoding proteins, the genome forms extensive RNA secondary structure, particularly in the 5*’*and 3*’*untranslated regions (12, 13). A number of these structures have essential roles in replication and translation, as well as in the virus’s interactions with its hosts. The genome adopts two alternative conformations, a linear form that supports translation and a circular form, formed by base pairing between the 5*’*and 3*’*ends, that is required for replication (14, 15). In the 3*’*untranslated region, exoribonuclease-resistant (xrRNA) elements fold into compact knots that stall the host exonuclease XRN1 partway through degradation, leaving a subgenomic flaviviral RNA (sfRNA) (16, 17, 18).

In Zika virus, this sfRNA antagonizes the type I interferon response (19) and is required for efficient transmission by mosquitoes (20), tying a noncoding RNA structure directly to immune evasion and vector competence. Other elements guide individual steps of replication, such as a capsid-region hairpin (cHP) that selects the translation start codon (21) and stem-loop A (SLA), which recruits the viral polymerase to initiate RNA synthesis (22).

These structural elements are distributed across the genome’s untranslated regions and capsid-coding sequence (Figure 1). At the 5*’*end they include SLA, stem-loop B (SLB), and cHP, together with the capsid-region pseudoknot DCS-PK (downstream of 5*’*cyclization sequence pseudoknot), which mediates the switch between the linear and circular forms (23, 24). The 3*’*untranslated region contains xrRNA1 and xrRNA2, two dumbbell structures (DB1 and DB2), a small hairpin (sHP), and the terminal 3*’*stem-loop (3*’*SL). In the circular form the upstream and downstream AUG regions (UAR and DAR), the cyclization sequence (CS), and a 5*’*–3*’*terminal stem base pair to join the two ends. We label the dumbbells DB1 and DB2 in 5*’*-to-3*’*order, even though this naming is reversed in parts of the literature (15, 23).

**Figure 1.**
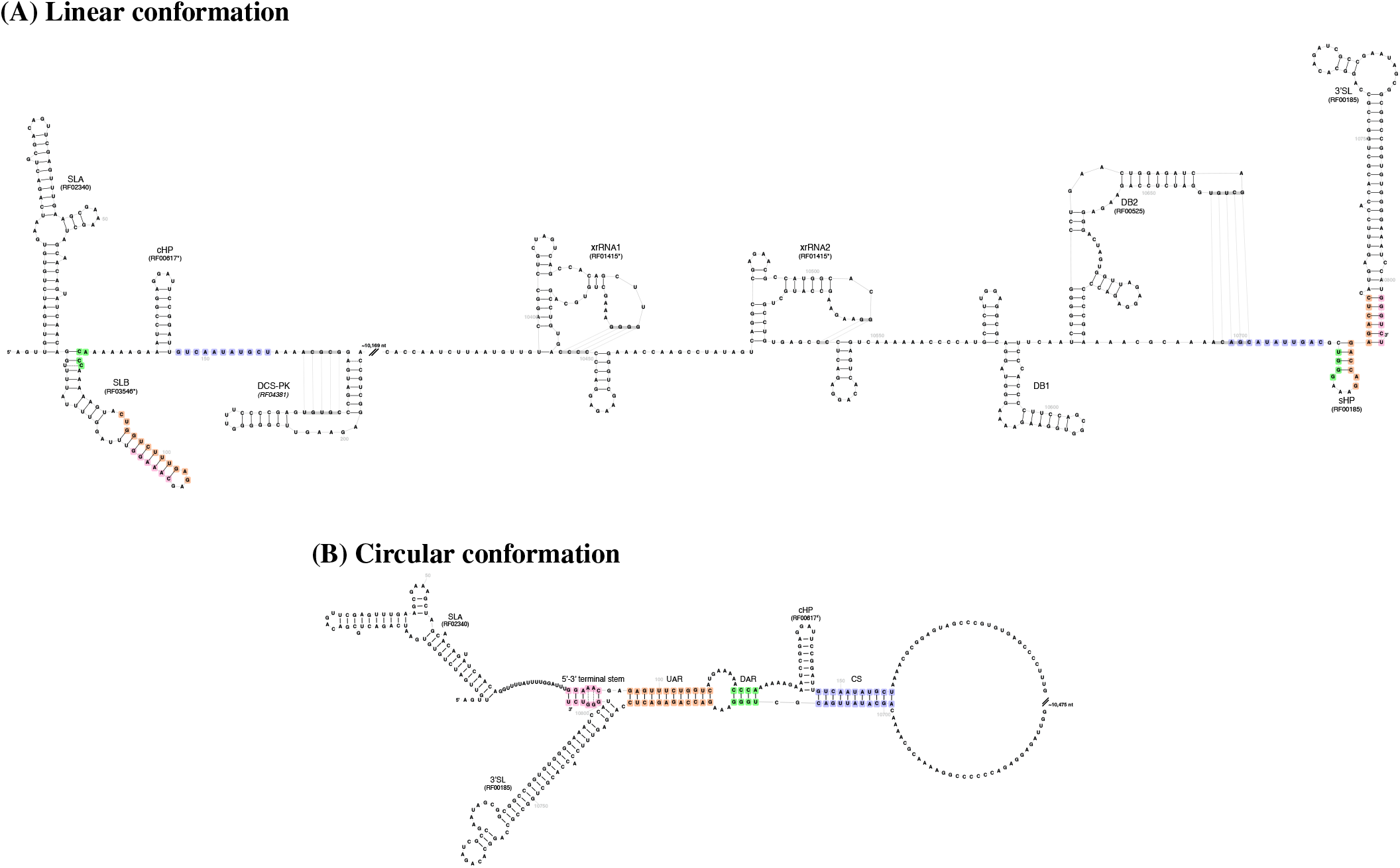
R2DT structural diagrams of the Zika virus 5’ /3’ untranslated and flanking structured regions, produced by `v-annotate.pl --draw r2dt` on NC 035889.1, the RefSeq genome that serves as one of the model’s reference sequences, using the two custom R2DT templates built for this project (zika-linear and zika-circular). Each template spans the structured 5 and 3 ends of the genome and omits the region between them, marked by a “//” break labeled with the number of omitted nucleotides. The two panels omit different amounts. The linear panel omits about 10,169 nucleotides of protein-coding sequence, and the circular panel omits about 10,475 nucleotides, which also includes xrRNA1, xrRNA2, DB1, and part of DB2. Nucleotides are drawn in black, and base-pairing is shown by connecting lines between paired residues. Two base pairs in each panel are drawn with dashed rather than solid lines. These pairs belong to the model’s consensus structure, but the residues that form them in NC 035889.1 are not Watson-Crick or G-U pairs, so R2DT’s drawing pipeline removes them before rendering. They are shown so that each panel displays the complete consensus structure. Genome coordinates are marked at multiples of 50, and the UAR, DAR, CS, and 5’ –3’ terminal cyclization stems are highlighted in orange, green, blue, and pink, on both panels. Element names, conformation labels and Rfam accessions, the highlighting of those four stems, and the two dashed base pairs were added after rendering and are not part of the native `--draw r2dt` output. The genome coordinates and the “//” break marker were produced with a locally modified R2DT when these panels were drawn. VADR v1.7.1 produces both natively, although its coordinate labels fall at regular intervals along the drawn structure rather than at multiples of 50. An accession in plain type indicates that the element matches its own Rfam family above that family’s gathering threshold (GA), an asterisk indicates a match to a related family below its gathering threshold, and italic type indicates the new family RF04381 created for this work. The circular panel labels only those elements that form in the cyclized conformation. **(A)** Linear (uncyclized) conformation, comprising in 5’*→* 3’ order SLA, SLB, cHP, the DCS-PK pseudoknot, xrRNA1, xrRNA2, the DB1/DB2 tandem dumbbells, sHP, and the 3’ SL. **(B)** Circular (genome-cyclized) conformation of the same regions, additionally showing the long-range 5’ –3’ base-pairing motifs that mediate genome cyclization. These are UAR, DAR, CS, and a 5’ –3’ terminal stem pairing the 5’ half of SLB with the 3’ terminal nucleotides. The linear and circular layouts were drawn to resemble Figure 1 of Li et al. (15).

### Zika data resources

Several databases support Zika research. The INSDC databases (GenBank, ENA, and DDBJ) are mirrored repositories that together host all publicly available Zika sequences, and NCBI Virus provides access to the GenBank holdings, letting users filter sequences by metadata such as collection date, geographic location, and genotype (25). The Bacterial and Viral Bioinformatics Resource Center (BV-BRC, formerly ViPR) adds integrated analysis tools (26), and its genome annotation service annotates viral genomes, including Zika virus, with VIGOR4, which predicts coding features by similarity to curated reference proteins (27). The European Virus Bioinformatics Center connects virologists and bioinformaticians and curates a catalog of virus bioinformatics tools and resources (28). The genetic diversity of Zika virus has been characterized phylogenetically, and here we follow the clade nomenclature of Seabra et al. (10). Despite these resources, Zika genomics still lacks standardized and curated reference sequences (29), and RNA secondary-structure information is largely missing from existing annotations, even though it is central to the viral life cycle.

### VADR is a general tool for viral genome annotation

VADR (Viral Annotation DefineR) is a software package for viral sequence annotation and validation used by the GenBank database for screening some types of incoming viral sequence submissions, and is available as a standalone portable software package for local use (30, 31). VADR is general in that it allows construction of a model for any viral species, from a reference single sequence or multiple sequence alignment. Any features for which there are reference coordinates can be included in the output annotation, including CDS, gene, mat_peptide, stem loop, and ncRNA features. VADR includes a script (v-annotate.pl) for validating and annotating input sequences using the reference model and features, outputting feature table files that can be sent to GenBank along with sequence submissions. The script also identifies dozens of types of unusual features and reports them as alerts in the output. Some alerts, like early stop codons in CDS regions or frameshifted regions, are fatal in that they cause a sequence to *fail*, which means the sequence should be inspected by a GenBank curator prior to its deposition into the database.

The Rfam database is a collection of RNA families, each represented by a multiple sequence alignment and a covariance model, that can be used to identify homologous structured RNAs in new sequences (32, 33). It includes families for many flavivirus RNA structural elements, but before this work it did not contain a family for the DCS-PK pseudoknot. The R2DT program (34) builds a template structure and alignment model and draws per-sequence secondary-structure diagrams, which is especially useful for flaviviruses like Zika virus that carry extensive structure in their 5*’*and 3*’*UTRs. The RNAcentral and Rfam databases together provide hundreds of such templates, some for flaviviruses, but not all of the Zika virus elements are represented.

In this work we address these gaps with a set of resources for Zika sequence annotation and analysis. At its core is a single alignment-based VADR model that annotates both the protein-coding features and the RNA secondary-structure elements of any Zika genome and produces GenBank-submission-ready feature tables together with the quality-control alerts used for submission screening. Alongside the model we provide R2DT templates that draw the structural elements of each sequence in its linear and circular conformations, a nearest-neighbor classifier that assigns sequences to Zika clades, and a new Rfam family for the DCS-PK pseudoknot (RF04381^1^). These resources are summarized in Table 1. Together these make secondary-structure information, which is largely absent from existing Zika annotation, available for every Zika sequence.

**Table 1.** Components of the Zika VADR resource.

| Component | Description | Location / accession |
| --- | --- | --- |
| Annotation + classification model package | Single covariance model (zika, 8-sequence training alignment) with protein, structural-RNA, and GenBank-QC annotation; nearest-neighbor clade classifier (African/Asiatic/American and Oceania), 100% agreement with Seabra et al. 2022); 97.2% pass rate on a 1254-sequence fresh test set (Table 4) | bitbucket.org/nawrockie/vadr-models-zika |
| R2DT structure templates | Linear (638-column) and circular (332-column) secondary-structure diagram templates spanning the 5'UTR, capsid region (incl. DCS-PK), and 3'UTR, rendered per-sequence by `v-annotate.pl --draw.r2dt`<sup>1</sup> | bitbucket.org/nawrockie/vadr-models-zika |
| DCS-PK Rfam family | New Rfam family for the flavivirus downstream-of-cyclization-sequence pseudoknot; 19 seed / 3 full sequences; GA/TC/NC 38.00/41.30/34.70 on Rfamseq 15.0 | Rfam accession RF04381 (Flavivirus_DCS-PK)<sup>2</sup> |
| Reproducibility package | Scripts and data underlying the benchmark, VIGOR4 comparison, Seabra confirmatory results, and the DCS-PK seed alignment | github.com/nawrockie/vadr-zika-paper-supplementary-material |
| R2DT diagram set | Secondary-structure diagrams for the test-set sequences, in the linear and circular conformations | same repository |

<sup>1</sup> Requires a VADR v1.7.1 installation with the `--draw.r2dt` flag and an R2DT installation with the Zika templates above installed as local data. <sup>2</sup> RF04381 will be included in the next release of Rfam after 15.10.

## MATERIALS AND METHODS

### Test and training sequence data sets

We created a dataset of freely available Zika virus sequences from GenBank with the following NCBI Entrez Direct (35) command: esearch -db nuccore -query “txid64320[Organism:exp]”| efetch-format fasta on May 4, 2026. From this set of 2725 sequences, we removed sequences with “UNVERIFIED” in their sequence description or belonging to the SYN (synthetic) or PAT (patent) divisions and any sequence less than 10,267 nt (95% of RefSeq NC 035889 length of 10,808) to create a full-length sequence dataset of 1262 sequences. We then selected and aligned the eight sequences listed in Table 2 to build our profile model. These eight sequences were selected because together they produce a model that accommodates the full sequence diversity of the 1262-sequence dataset while allowing our nearest-neighbor based classification strategy to reproduce the taxonomic group assignments of all sequences in a published study exactly (10). The remaining 1254 sequences formed our test set.

**Table 2.** The eight Zika virus sequences comprising the model’s reference training alignment, listed by clade and collection year. Clades follow the three-way nomenclature used throughout: ZA (African), ZB.1 (Asiatic) and ZB.2 (American and Oceania). Country, collection year and length are from the GenBank records.

| Accession | Clade | Country | Year | Length (nt) |
| --- | --- | --- | --- | --- |
| KY989511.1 | ZA (African) | Uganda | 1947 | 10,807 |
| KX377336.1 | ZB.1 (Asiatic) | Malaysia | 1966 | 10,807 |
| MT377504.1 | ZB.1 (Asiatic) | Thailand | 2006 | 10,603 |
| MG807646.1 | ZB.1 (Asiatic) | Thailand | 2016 | 10,807 |
| MW915411.1 | ZB.1 (Asiatic) | Viet Nam | 2016 | 10,661 |
| MN611472.1 | ZB.1 (Asiatic) | China | 2019 | 10,807 |
| PQ551088.1 | ZB.1 (Asiatic) | Kenya | 2024 | 10,666 |
| NC.035889.1 | ZB.2 (American and Oceania) | Brazil | 2015 | 10,808 |

NC.035889.1 is the Zika virus RefSeq. PQ551088.1 (isolate AFI-LAM-1993) is a 2024 case detected in Kenya that belongs phylogenetically to the Asiatic clade (ZB.1) rather than to the African lineage, a reminder that clade membership here is phylogenetic and not geographic. It was added to the training set to represent the original, endemic part of ZB.1, which the other ZB.1 training sequences do not cover, and to resolve the four boundary cases MK238035–MK238038 discussed below.

### Reference annotation of coding and noncoding features in Zika genome

Our Zika model includes one CDS feature (polyprotein), one gene feature, 14 mat peptide features, one ncRNA feature, and ten structural RNA stem loop features. The annotation of the CDS, gene and mat peptide features derives from the GenBank annotation of the RefSeq NC 035889.1 sequence, which is one of the eight sequences in our model’s training alignment. For the stem loop structures, we used Infernal v1.1.5 and Rfam v14.10 to identify plausible Rfam matches in the genome and truncated them to the structured stem loop regions (not including any terminal single stranded nucleotides). The lone ncRNA feature is the subgenomic flavivirus RNA (sfRNA1), a 3*’*UTR-derived noncoding RNA produced by host exonuclease XRN1 stalling at the xrRNA1 structure; we annotate it following the convention used in VADR’s dengue models. We compared and, where necessary, revised these annotations against published reports on these elements (15, 17, 22, 23, 36, 37, 38, 39).

### Covariance model construction

Our eight sequence profile alignment was created by first generating a predicted secondary structure for the NC 035889 sequence based on the structures of the stem loop features from the Infernal/Rfam predictions, creating a single sequence Stockholm ‘alignment’, building a covariance model (CM) from that (Infernal v1.1.5 `cmbuild` program) and aligning the eight training sequences to that model (`cmalign -O`). We then ran `cmbuild` using that alignment as input to create the final model (`cmbuild -F --ere 1.1 --p7ml --hand --fraggiven -n zika`), with `--hand` specifying consensus columns from the reference annotation, which corresponds to the 10,808 positions of RefSeq NC 035889.1, `--fraggiven` taking fragment definitions from the input alignment, in which seven of the eight training sequences are marked as fragments (four of them lacking only a single terminal nucleotide, and the remaining three lacking 142, 147 and 205 nucleotides at one or both ends), `--p7ml` building the maximum-likelihood profile-HMM filter, and `--ere 1.1` setting the target relative entropy to 1.1 bits, a value chosen by an `--ere` sweep because it correctly classifies a one-nucleotide 5*’*-truncation case. We also added taxonomic group and subgroup annotation, an alternative secondary structure annotation for the circular conformation (`SS cons2`) derived from (15), and a tertiary structure annotation (`SS cons tertiary`) recording three non-canonical pairs within xrRNA1 derived from (18), to the eight sequence alignment to use with `v-annotate.pl`.

To assess whether a single profile model matches the more common one-model-per-group approach, we also built a library of three single-sequence models, one per clade, from RefSeqs KY989511 (ZA, African), MG807646 (ZB.1, Asiatic), and NC 035889 (ZB.2, American and Oceania), each annotated with the same feature set as the profile model. We compared the two libraries on the full test set in terms of per-sequence pass/fail agreement, per-feature coordinate identity, clade classification, and run time. Run times were measured on a 50-sequence clade-stratified subset, with both libraries run single-threaded and back-to-back on a dedicated compute node with an Intel Xeon Gold 5415+ processor (2.9 GHz).

### DCS-PK pseudoknot characterization and Rfam submission

The DCS-PK RNA secondary structure element is a functional element that is not represented by an Rfam model, so we built an alignment and created an Rfam family (RF04381) which will appear in the next release of Rfam. Starting from the published predicted secondary structure of the Zika DCS-PK element (37), we built a single-sequence Stockholm alignment, constructed a CM from it, and searched a large viral sequence database derived from GenBank in August 2025 including 3.86 million viral sequences totalling approximately 14.7 Gb. We iteratively expanded the seed by adding representative flavivirus homologs recovered from this search (including Kokobera, Baiyangdian (duck Tembusu), Murray Valley encephalitis, Usutu, T’Ho, and Ilheus viruses) and correcting two mislabeled database sequences that were in fact flaviviruses, realigning at each step. We then used R-scape (version 2.5.9.c) in its “evaluate given structure” mode (`-s`) to assess covariation support for the proposed base pairs. The final seed alignment was submitted to Rfam and searched against Rfamseq 15.0 to set the family’s gathering (38.0 bits), trusted-cutoff, and noise-cutoff thresholds (GA, TC, and NC respectively); RF04381 was assigned to Rfam clan CL00129 alongside the related cHP (RF00617), DENV SLA (RF02340), and Flavivirus-5UTR (RF03546) families.

### VADR annotation recommended settings

We ran VADR (version 1.7.1) and experimented with the use of various command-line flags with `v-annotate.pl`, settling on the following recommended usage:

~~~
  v-annotate.pl -r --r file zika.rpn.fa
--nosub
--lowsim5ftr 15 --lowsim3ftr 15
--draw r2dt
~~~

The flags above are those we recommend; the model files themselves are supplied separately. The Zika models are distributed as the `vadr-models-zika` package (Table 1), and are selected by passing the unpacked package directory to `--mdir` together with `--mkey zika`.

`v-annotate.pl` uses `blastn` (40, 41) to identify the closest reference sequence for N-replacement and `blastx` to validate the protein translation of each predicted coding sequence against a set of reference proteins.

Here, `-r` enables N-replacement, in which runs of ambiguous nucleotides in an input sequence are temporarily replaced with nucleotides from the closest matching reference sequence prior to annotation, to prevent alerts due solely to the Ns (the original Ns would be replaced prior to deposition in GenBank) (31). The `--r file` option supplies a database of the eight training genomes for this step (instead of the default single consensus), improving fill fidelity by doctoring from a same-clade sequence. `--nosub` uses the alignment program’s (`cmalign`’s) default handling of truncated sequences rather than VADR’s default sub mode, which predicts the model start and end positions first and aligns to a sub-CM spanning only that range. Sub mode can misplace a residue at a truncated 5*’*boundary and raise spurious alerts. `--lowsim5ftr 15` and `--lowsim3ftr 15` raise the minimum length of a low-similarity region at a feature’s 5*’*or 3*’*end needed to trigger a low-similarity alert to 15nt, which prevents a few failures we observed during testing due to short benign terminal substitutions. The `--draw r2dt` flag adds a post-processing step that runs the R2DT program (34) using a Zika-specific RNA secondary structure template to generate a secondary structure image file of the input sequence folded into the two alternative conserved secondary structures (including four pseudoknots), consisting of mainly the 5*’*and 3*’*ends of the Zika genome. The linear structure includes 192 basepairs, 118 of which are shared with the circular structure; the circular structure has 33 basepairs of its own. These structure template layouts were drawn to resemble Figure 1 of Li et al. (15) using the RNAcanvas structure editing tool (42).

### Classification strategy and testing

We use the nearest neighbor-based classification strategy implemented in `v-annotate.pl` to classify each input Zika sequence *S* into one of three clades, ZA (African), ZB.1 (Asiatic), or ZB.2 (American and Oceania), by computing the percent identity between *S* and all eight training sequences (as aligned in the model’s reference alignment) and defining *S*’s clade as the clade of most similar training sequence.

To test the ability of our model to accurately classify Zika sequences into these three clades, we used the 760 sequence dataset from the Seabra et al. study (10). Seabra et al. proposed a hierarchical nomenclature comprising two main lineages, ZA and ZB, corresponding to the African and Asian genotypes; ZB is divided into ZB.1 and ZB.2, and each of these is further subdivided into three sublineages (ZB.1.0– ZB.1.2 and ZB.2.0–ZB.2.2). Our classifier distinguishes sequences at the coarser ZA/ZB.1/ZB.2 level, and we therefore assess agreement with Seabra et al.’s assignments at that level.

### Comparing with VIGOR4

To compare VADR’s ability to annotate Zika sequences with another freely available and widely used viral annotation tool, VIGOR4, we selected the 981 of our 1254 test-set sequences that contained no run of 20 or more consecutive ambiguous nucleotides, and compared the `v-annotate.pl` output annotations with VIGOR4-generated annotations in terms of the coordinate spans of features. We excluded sequences with a run of *≥* 20 ambiguous nucleotides because VIGOR4 treats any such run as a sequencing gap (its `min seq gap length` default) and splits the gene model spanning it into independently scored fragments, roughly halving VIGOR4’s reported reference coverage; sequences without Ns or with scattered Ns but no such runs annotate essentially identically between the two tools.

VIGOR4 (v4.1) was run as `vigor4 -i input.fa -o output -d zikv db --overwrite-output`. VIGOR4 feature coordinates were taken from its `.pep` output (whose FASTA headers give the CDS and mat peptide locations) and compared to the corresponding VADR `.ftr` output for coordinate identity. VADR reports each sequence as passing or failing, whereas VIGOR4 does not assign a pass/fail status or emit alerts; for the purpose of our comparison we considered a sequence to fail VIGOR4 when VIGOR4 produced no gene model (no annotated CDS) for that sequence.

## RESULTS

This work has produced computational resources for Zika sequence validation, annotation and analysis (Table 1). These resources are all freely available to use with the VADR and R2DT software packages and in the next release of the Rfam database.

Many VADR models are based on single sequences, including the dengue virus, which has four single-sequence models, one per serotype. We opted to build one model based on an alignment for Zika to make maintenance simpler (it is easier to keep one model up to date than multiple models), because the new nearest-neighbor method allows us to classify sequences into multiple groups without requiring one model per group, and because our tests showed nearly identical performance of the single profile model versus multiple single-sequence models. On the test set of 1254 nearly full length sequences, the pass/fail designations were 1219 pass and 35 fail (97.21%), and on the set of 1026 partial sequences they were 1013 pass and 13 fail (98.73%). On the full test set, the single profile model and the three-single-sequence-model library agreed on the pass/fail verdict for 99.3% of sequences and on feature coordinates for 99.96% of shared annotations, and on clade for 97.5% (all disagreements falling at the closely related Asiatic and American clades); the profile model ran about 5% faster.

Thirty-five of the 1254 full-length test sequences were classified as fails by `v-annotate.pl` (Table 3). The largest category (19) carries a high-confidence possible-frameshift alert (`fsthicfi`): in every case a deletion and a nearby insertion restore the reading frame before the end of the CDS, so the polyprotein remains full-length (all are GenBank “complete cds” records) and no frame-disabling alert occurs anywhere in the benchmark. Five sequences carry premature CDS stop codons (`cdsstopn`/`cdsstopp`, the same in-frame stop caught independently by nucleotide- and protein-based alignment); one more, MW123924, has a mutated or absent expected stop codon and is flagged at the first in-frame stop further downstream (`mutendcd`/`mutendex`) rather than a premature one; and the remaining ten fail on 5*’*/3*’*feature-boundary or low-similarity issues, one of which, MW123925, fails on a low-confidence 5*’*boundary in the structured SLA element, where it carries an off-consensus nucleotide. We consider all 35 warranted. Each flags a genuine sequence anomaly a GenBank curator would want to inspect before automatic deposition (though many, such as the compensated frameshifts, would pass on inspection).

**Table 3.**
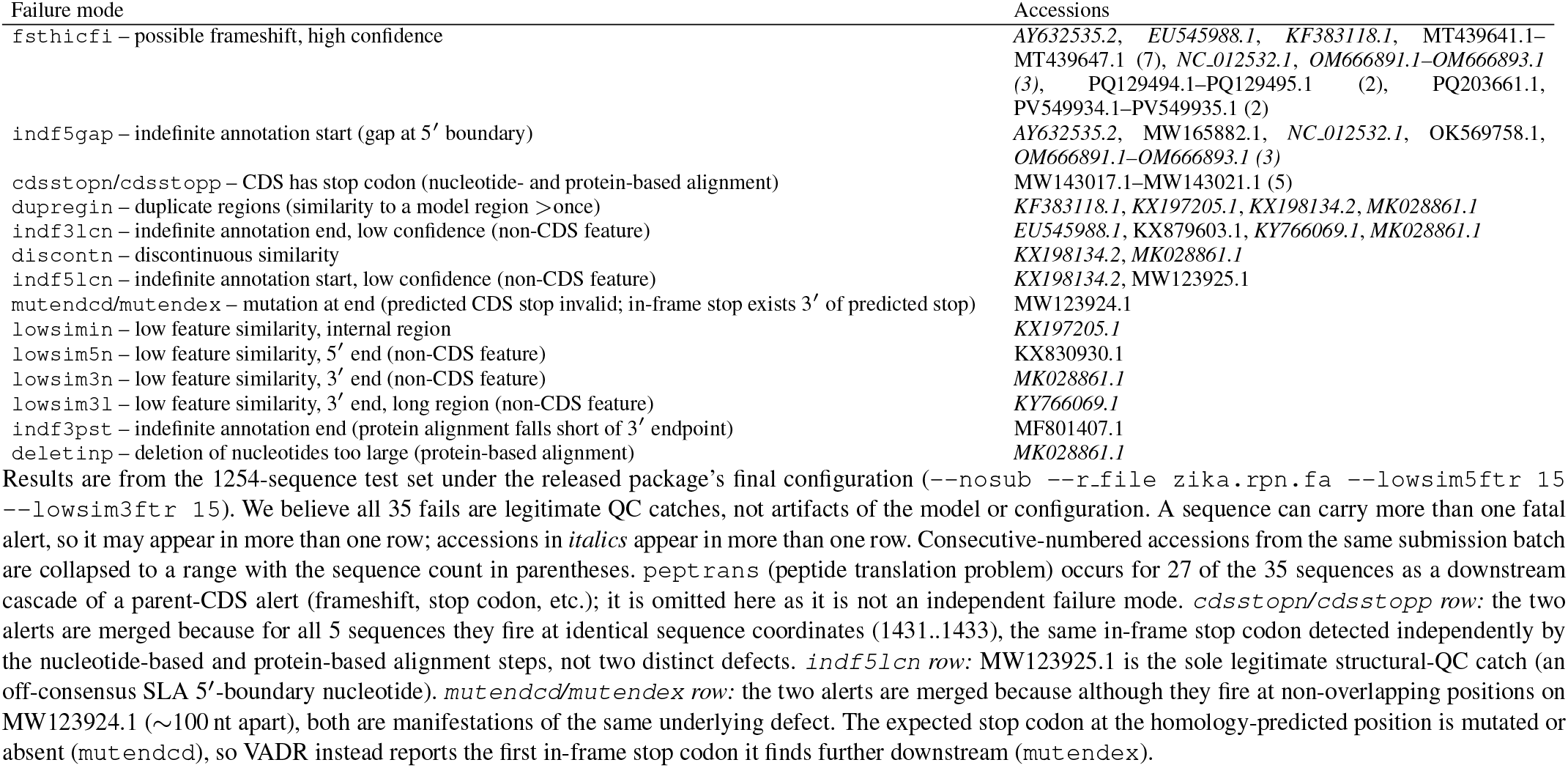
Fatal alert codes for the 35 full-length test-set sequences (of 1254) failing under the released package’s final configuration. Frameshifts (fsthicfi) and CDS stop codons (cdsstopn/cdsstopp) dominate the primary causes. peptrans is omitted (see footnote).

Of the 1026 partial sequences, 13 failed. VADR annotated the covered region correctly in all 13. Five carry genuine defects that would fail at any length, including large deletions and an internal indel that splits the protein alignment. Seven are correctly annotated legitimate partial sequences that would pass were they full length, failing only because a terminal coding boundary falls short of the expected endpoint or because coverage drops below the threshold on a very short fragment. The remaining sequence, AY326412, is borderline, a 3*’*UTR-only fragment whose coverage is reduced by a genuine internal region of low similarity.

We compared `v-annotate.pl` and VIGOR4 annotations and found that the overwhelming majority agree: across the 981 sequences, 11,709 of 11,712 shared features (CDS and mat peptide) had identical coordinates, and 974 of the 981 sequences (99.3%) were fully concordant. Of the remaining seven, five are the block described below for which VIGOR4 produced no gene models at all, and two carry coordinate mismatches: three mismatched features in total, all boundary shifts of at most three nucleotides. The VADR model additionally annotates features absent from the VIGOR4 model, comprising three mat peptides (capsid protein C, anchored capsid protein C, and membrane glycoprotein precursor M), ten stem loop features, and one ncRNA feature, which could not be compared. Across the 981 sequences this adds about 12 annotated features per sequence that VIGOR4 does not produce (12,168 in total). Unlike VADR, VIGOR4 does not assign a pass/fail status or emit alerts, and so flags none of the sequences VADR fails on quality grounds; the one exception is a contiguous block of five sequences (MW143017.1–MW143021.1) carrying genuine in-frame stop codons, for which VIGOR4 produced no gene models and VADR likewise reports failures. VADR’s quality-control alerts thus provide submission screening that VIGOR4’s annotation-only output does not.

The nearest-neighbor classification strategy showed perfect agreement (754 of 754 classified sequences, 100%) with the Seabra et al. clade assignments (10) at the ZA/ZB.1/ZB.2 level, collapsing Seabra’s finer sublineages (Table 4). We note that our eight training sequences were selected in part because they reproduced these classifications.

**Table 4.** Clade-classification agreement and nearest-neighbor confidence, by clade, against the Seabra et al. (2022) reference assignments (754 non-training test sequences).

| Clade | N | Agreement | Mean %id to best | Margin to 2nd, pp (median, IQR) | Min | Max |
| --- | --- | --- | --- | --- | --- | --- |
| ZA (African)<sup>1,2</sup> | 7 | 7/7 (100%) | 95.2% | 4.85 (3.54–4.89) | 3.11 | 9.41 |
| ZB.1 (Asiatic) | 101 | 101/101 (100%) | 99.4% | 0.69 (0.54–0.70) | 0.01 | 2.62 |
| ZB.2 (American and Oceania) | 646 | 646/646 (100%) | 99.4% | 0.49 (0.47–0.51) | 0.10 | 0.88 |
| Total | 754 | 754/754 (100%) | 99.4% | – | – | – |

<sup>1</sup> Clade classification is by nearest-neighbor % identity to the model’s training sequences, as reported in *v-annotate.pl*’s classification output (the *.scn* file). Mean % id is the identity to the nearest training sequence. Margin is that identity minus the identity to the nearest training sequence of a different clade, in percentage points. Six of the eight training sequences (Table 2) are present in the 760-sequence set and are excluded as queries, leaving 754. The Total row omits pooled margin statistics because the per-clade margin distributions are bimodal, the African tail against the tight Asiatic/American cluster, making a pooled median and IQR misleading. <sup>2</sup> Agreement is against the Seabra et al. (2022) reference clade assignments.

This assignment is nonetheless reported as indefinite for almost every sequence. VADR emits an indefinite-classification alert (`nnindfcl`) when a sequence’s margin to the next-best clade falls below a threshold, 5 percentage points by default, and 753 of the 754 sequences (99.9%) fall below it. Every Asiatic and American sequence does, with median margins of only 0.69 and 0.49 percentage points (Table 4), as do six of the seven African sequences despite their considerably wider margins; the sole exception is AY632535.2, at 9.41 percentage points. The alert is never fatal and changes neither the annotation nor the pass/fail status, and the clade assignments were fully concordant despite nearly all of them carrying it. It therefore has little diagnostic value for Zika virus, whose clades are separated by far less divergence than the default threshold anticipates. The threshold can be adjusted with the `--nn indefclass` option.

Secondary structure elements in flaviviruses including Zika have been analyzed via comparative sequence analysis and experimental techniques (13, 14, 15, 22, 36, 37, 39, 43, 44, 45). Our VADR model incorporates ten base-paired stem loop structural elements. Searching the Zika genome against all Rfam families with Infernal (33), and retaining hits with E-value below 1 after discarding two taxonomically implausible matches to human long noncoding RNAs (RF01954, SOX2OT, *E* = 0.55; RF02204, WT1-AS, *E* = 0.93), recovered eight of the ten: SLA, DB2, sHP, and the 3*’*SL matched their Rfam families (RF02340, RF00525, and RF00185) above the family gathering threshold, while SLB, cHP, xrRNA1, and xrRNA2 matched related families (RF03546, RF00617, and RF01415) below it. Only DB1 and DCS-PK were not detected, for different reasons. DB1 is a degenerate dumbbell (ΨDB) too diverged to be recognized by the flavivirus dumbbell family (RF00525) that DB2 matches (23, 39). DCS-PK, by contrast, is not modeled by any Rfam family, so we created an Rfam family (RF04381) of length 51, containing 19 sequences from 14 flavivirus species (dengue, Zika, West Nile, Japanese encephalitis, Murray Valley encephalitis, Usutu, St. Louis encephalitis, Spondweni, Kokobera, Cacipacore, Ilomantsi, Baiyangdian, T’Ho, and Ilheus viruses)^2^ and comprising 18 base pairs, of which five form the pseudoknot. R-scape analysis (46) shows that ten of the 18 base pairs significantly covary across the 19 seed sequences, supporting the experimental evidence (37, 44) that these base pairs are functionally relevant and conserved. Figure 2 shows the structure, conservation, and covariation of the RF04381 seed alignment.

**Figure 2.**
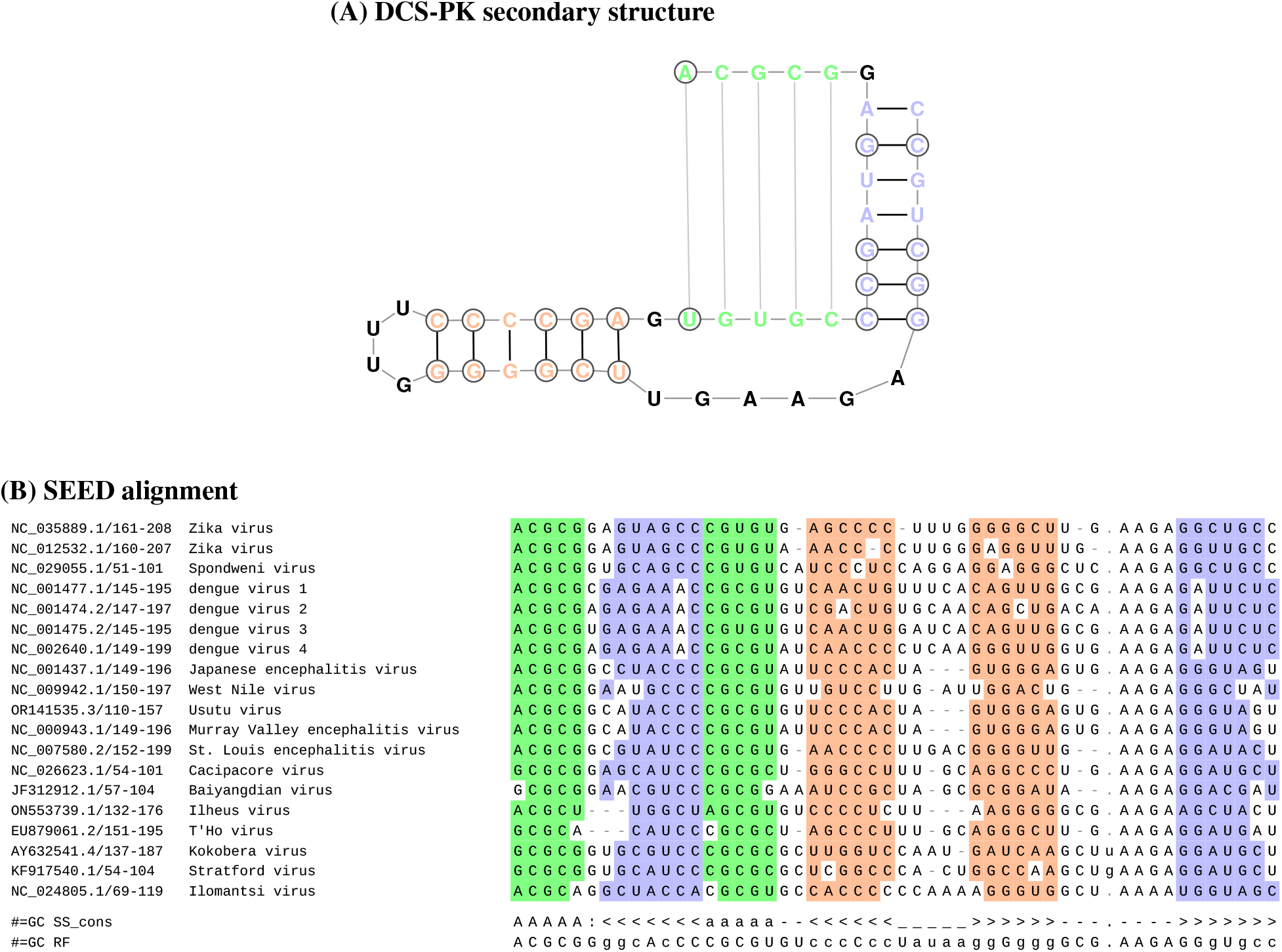
The DCS-PK (Downstream of 5 Cyclization Sequence PseudoKnot) element, submitted to Rfam as family `RF04381` (`Flavivirus DCS-PK`; clan `CL00129`, alongside cHP, DENV SLA, and Flavivirus-5UTR, resolving a 2-nt boundary overlap with `RF03546` (Flavivirus-5UTR)) and added to pending families in July 2026 for the next release of Rfam. A search of Rfamseq 15.0 using the family’s gathering threshold (GA 38.00 bits; TC 41.30, NC 34.70) identified three sequences above GA, which constitute the family’s full alignment. These are dengue virus 1 `U88536.1` (58.7 bits), *Levilactobacillus bambusae* `QCXQ01000013.1` (50.8 bits, likely flavivirus contamination in a bacterial WGS assembly), and Zika virus MR 766 `AY632535.2` (44.6 bits). None of the 19 SEED sequences is part of the Rfamseq 15.0 sequence set searched, so the seed and full sets are disjoint, and RF04381 comprises 19 seed and 3 full sequences. **(A)** Secondary structure of the DCS-PK pseudoknot in Zika virus NC 035889.1, drawn as in Figure 1 and colored by structural element as in panel B (pseudoknot green, stem 1 blue, stem 2 orange; unpaired residues black). Across the 19-sequence SEED alignment, ten of the 18 consensus base pairs covary significantly by R-scape in `-s` (“evaluate given structure”) mode (*E* < 0.05, against an expectation of 3.1 *±* 1.5 by chance; the weakest of the ten falls just under the threshold at *E* = 0.045), with stem 2 the most strongly supported helix; the 20 residues forming these ten pairs are circled. **(B)** The 19-sequence SEED alignment, with the consensus secondary structure (`SS cons`, including the pseudoknot in `A`/`a` notation) and the reference annotation (`RF`) shown below it. At each base-paired position a residue is colored by its structural element (pseudoknot green, stem 1 blue, stem 2 orange) when it forms a Watson-Crick or G-U pair with its partner, and is left white otherwise.

The linear and circular conformations differ in which structural elements fold. In the linear form, all ten elements fold locally: SLA, SLB, and cHP in the 5*’*region; the DCS-PK pseudoknot in the capsid-coding region; and xrRNA1, xrRNA2, DB1, DB2, sHP, and the 3*’*SL in the 3*’*UTR. In the circular form, the 5*’*and 3*’*ends instead base pair directly through four long-range cyclization stems (the UAR, DAR, and CS stems, plus a 5*’*–3*’*terminal stem, which has no standard name in the literature, pairing the 5*’*half of SLB with the 3*’*terminal nucleotides; 33 base pairs in total). This refolding displaces the DCS-PK pseudoknot, which forms only in the linear conformation (15), and draws the 3*’*half of SLB into the cyclization interaction (14); the remaining elements are retained.

## DISCUSSION

The VADR profile model reported here provides a rich source of sequence and secondary structure information for Zika researchers. Because sequence databases often lack structural information, the model’s secondary structure annotations are of particular value. They are delivered for entire genomes and in multiple formats: sequence-coordinate annotations of structural elements, structure-annotated alignments, and per-sequence secondary structure diagrams of any Zika sequence. They also include both the linear and circular conformations of the genome, making it easier to visualize how individual nucleotide mutations affect each structural element.

The DCS-PK element is central to the conformational switch between the linear and circular conformations of the genome: it folds locally in the linear form and unfolds during cyclization, allowing the 5*’*and 3*’*ends to base pair (15, 23). In Zika, DCS-PK has adapted over the virus’s evolutionary history. Its structural stability has increased, and the more stable Asian-lineage element supports more efficient replication than the early-diverged African-lineage version (24). With this work, DCS-PK has been added to the Rfam database as family RF04381 and will be incorporated into the annotations of other tools that use Rfam data, enabling its detection across mosquito-borne flaviviruses.

The profile model developed here produces annotations and taxonomic classifications nearly identical to those of a library of individual single-sequence models, agreeing on 1245 of 1254 pass/fail verdicts (99.3%) and 1222 of 1254 clade assignments (97.5%). We prefer the single model because it separates classification from annotation, so a sequence near a clade boundary is annotated against the same reference model no matter which clade it is assigned to. In a per-clade library the annotating model changes with the classification, and as the margins described below show, those calls can turn on a few tenths of a percentage point of identity. One model also keeps every sequence in a single coordinate frame, and a newly recognized lineage can be accommodated by adding sequences to the alignment rather than by building, calibrating and validating another model. Its annotations are likewise nearly identical to those of VIGOR4 (974 of 981 sequences, 99.3% concordant), but our model additionally annotates three mature peptides, ten structural RNA stem-loops, and one ncRNA feature that VIGOR4 does not.

The taxonomic classification ability of our model showed perfect agreement with the Seabra et al. classification (10), but to be fair, we trained our model to perform well versus that dataset. The per-clade calls are made based on percent identity to each of our eight training sequences in an alignment, and the margins are often small. For example, four sequences (MK238035–MK238038) from a single study (47) sit on the Asiatic–American boundary; our classifier assigns them to the correct (Asiatic) clade, but by a margin of only about 0.3 percentage points of identity, and only after we added a training sequence specifically to resolve them (the main change we made to bring our training set into full agreement with that classification). Nevertheless, the model does provide an accurate way of classifying Zika sequences which we expect will be of utility to users. If and when future lineages emerge, it will be necessary to update our model to include sequences from those lineages, at which point testing will be required to ensure accuracy.

### Limitations and future directions

Some valid Zika sequences are flagged as failures by our model: 35 of the 1254 full-length test sequences (2.8%) and 13 of the 1026 partial sequences (1.3%). Of these 48 failures, 26 are sequences that VADR flags conservatively and that a curator would likely clear, ten flag genuine defects such as premature stop codons and large deletions, and the remaining twelve reflect terminal boundary, low-similarity, or stop-codon-position issues, with one partial sequence borderline on coverage. The 26 comprise the 19 full-length failures carrying compensated frameshifts that leave the polyprotein intact and seven partial failures that would pass were they full length. This behavior is a deliberate choice: our priority with VADR is to minimize false negatives, accepting some false positives in exchange, rather than the reverse. We have not experimentally verified that our annotations are correct or that every VADR-passing sequence is valid; the near-complete concordance with VIGOR4’s independent annotations is our principal external check, alongside the model’s agreement with published clade assignments.

A major limitation of VADR is its slow speed. Processing a single full length Zika sequence takes roughly 20 seconds. Processing can be easily parallelized across multiple cores via multithreading due to the independence of every sequence’s validation and annotation, and the number of Zika sequences (about 3400 in GenBank, requiring roughly 19 total CPU hours to process all of them) reduces the impact of this slow processing speed, but if sequencing ramped up it could become a limiting factor. VADR also has steep memory requirements and requires about 4 GB per core. As VADR is applied to more and more viral models, reducing this computational footprint will be an important aim.

The main goal for VADR development and use is to extend to more viruses. This work on Zika, combined with earlier work on dengue virus (30), another flavivirus that uses earlier VADR conventions (single sequence based models, no R2DT diagrams), demonstrates that VADR is a general tool for flavivirus sequence and structure analysis, and opens the door to the development of additional flavivirus models. Many flaviviruses share homologous RNA structures and some adopt cyclization and linear conformations similar to those of Zika. More broadly, pairing sequence validation with per-sequence secondary-structure annotation is a general strategy. It should apply to any virus whose genome carries conserved RNA structure.

## DATA AVAILABILITY

VADR and the Zika models are in the public domain and are freely available at https://github.com/ncbi/vadr and https://bitbucket.org/nawrockie/vadr-models-zika. VADR depends on the following software, which is downloaded and installed as part of VADR installation: Bio-Easel v0.18, BLAST+ v2.17.0, Infernal v1.1.5, FASTA v36.3.8h, minimap2 v2.30, R2DT v2.3 and Sequip v0.11. Instructions for using VADR for zika annotation can be found at https://github.com/ncbi/vadr/wiki/Zika-virus-annotation. The supplementary material is available at https://github.com/nawrockie/vadr-zika-paper-supplementary-material. It contains the scripts and data underlying the benchmark results, the VIGOR4 comparison, the Seabra confirmatory analyses, and the DCS-PK seed alignment, together with R2DT secondary-structure diagrams for the test-set sequences, each with a README describing its contents.

## ACKNOWLEDGEMENTS

We thank Alvin Crespo Bellido for assisting in tree-building to aide our classification efforts and Ron Patterson and his team for excellent support and management of NLM’s shared computing resources. The authors wrote and revised this manuscript, using a large language model (Claude, Anthropic) for assistance with drafting and editing text, code development, and benchmark analysis. The authors designed and directed the work, verified all results, and take full responsibility for the content.

## Footnotes

1 RF04381 is currently a pending Rfam family and will be included in the first Rfam release after 15.10.

2 Species assignments follow ICTV MSL40v2 (2024). The four dengue serotypes are one species (*Orthoflavivirus denguei*), as are Kokobera virus and Stratford virus (*Orthoflavivirus kokoberaorum*), both of which are represented in the seed alignment.

